# Age-dependent decrease in inhibitory drive on the excitatory superficial spinal dorsal horn neurons

**DOI:** 10.1101/2023.06.28.546829

**Authors:** Prudhvi Raj Rayi, Shaya Lev, Alexander M Binshtok

## Abstract

The excitatory and inhibitory interneurons of superficial laminae I-II of the spinal dorsal horn (SDH) receive and process pain-related information from the primary afferents and transmit it to the brain via the projection neurons. Thus, the interaction between excitatory and inhibitory SDH interneurons is crucial in determining the output from the spinal cord network. Disruption of this interaction in pathological conditions leads to increased SDH output to the higher brain centers, which could underlie pathological pain. Here, we examined whether the changes in the intrinsic SDH connectivity also occur with age, possibly underlying age-related increase in pain sensitivity. Using *Vgat;tdTomato* transgenic mouse line, we compared the spontaneous inhibitory postsynaptic currents (sIPSCs) in inhibitory tdTomato^+^ and excitatory tdTomato^−^ interneurons between adult (3-5 m.o.) and aged (12-13 m.o.) mice. We demonstrate that in adult mice, the amplitude and frequency of the sIPSCs in the excitatory interneurons were significantly higher than in inhibitory interneurons. These differences were annulled in aged mice. Further, we show that in aged mice, excitatory neurons receive less inhibition than in adult mice. This could lead to overall disinhibition of the SDH network, which might underlie increased pain perception among the aged population.

## 1. Introduction

The information encoding noxious stimuli from nociceptive peripheral afferents is processed by inhibitory and excitatory interneurons of the superficial spinal dorsal horn (SDH) before relaying it to the projection neurons [1]. A well-tuned interaction between these neurons is therefore required for normal nociception and pain. Indeed, in pathological conditions such as inflammation and nerve injury, the decrease in the inhibitory drive leading to the alteration of excitation-inhibition balance towards stronger excitation was demonstrated [2–4]. These studies emphasize the critical role of disinhibition in abnormal processing and relay of nociceptive signals to higher brain regions. Here, we asked whether the inhibitory synaptic transmission in SDH interneurons is altered with age, contributing to the age-related increase in pain. Human and animal studies have shown an increase in response to noxious stimuli with age [5]. Importantly, aging increases the susceptibility to chronic pain, causing suffering, disability, and social isolation [6]. It was suggested that a decrease in endogenous analgesic mechanisms, for example, loss of serotonergic and noradrenergic neurons in lamina I of the SDH, and impaired descending pain modulation, could underlie the age-related increase in pain sensation and susceptibility to pain chronification [7–9]. Using patch-in-slice recordings from genetically identified inhibitory and excitatory SDH interneurons, we showed that in adult mice inhibition on excitatory neurons is stronger than on inhibitory neurons. This inhibitory drive on excitatory neurons is altered with age, such that the excitatory interneurons in aged mice receive less inhibition than in the adults.

## 2. Materials and methods

### 2.1 Animals

The *Vgat;tdTomato* mice were generated by crossing Vgat-ires-cre knock-in line B6J.129S6(FVB)-Slc32a1^*tm2(cre)Lowl*^*/*MwarJ (JAX stock #028862) with Ai14 reporter mouse line B6.Cg-*Gt(ROSA)26Sor*^*tm14(CAG-TdTomato)Hze*^*/J* (JAX stock #007914). The resulting mice expressed tdTomato fluorescence selectively in neurons expressing vesicular GABA transporter (*Vgat*). Same-sex littermates were group-housed on a 12 h light/dark cycle with food and water *ad libitum*. Adult (3-5 m.o.) male mice were used for validating the firing properties of neurons expressing tdTomato (tdTomato^+^) and neurons with no expression of tdTomato (tdTomato^−^) from the laminae I-II of the superficial spinal dorsal horn (SDH). Adult male and female mice (3-5 m.o.) and old male and female mice (12-13 m.o.) were used for all voltage-clamp recordings. To assess the presence of the main effects of sex or interactions of sex with the properties of the synaptic transmission of the examined neurons, all data sets were analyzed using sex as a factor. We did not find any effects of sex or interactions, so the data were pooled for all reported analyses. All procedures were conducted in accordance with the guidelines and approval of the animal ethics committee of the Hebrew University of Jerusalem (MD18-15604).

### 2.2 Spinal cord slice preparation

Mice were anesthetized with a lethal dose of ketamine/domitor anesthetic mix (100 mg/kg i.p) and checked for any reflex by a toe pinch to ensure proper anesthesia. Following the development of deep anesthesia, mice were perfused with 20 mL ice-cold sucrose-based cutting solution containing (in mM): 110 sucrose, 5 D-glucose, 60 NaCl, 28 NaHCO_3_, 3 KCl, 1.25 NaH_2_PO_4_, 7 MgCl_2,_ and 0.5 CaCl_2_. The spinal cord containing lumbar enlargement was dissected from the ventral side to minimize the damage to the dorsal horns and glued the ventral side to an agar block with a slot to support the tissue while cutting. 300 μm transverse sections were cut using a Leica VT1200S vibrating blade microtome (Leica Biosystems), and the spinal cord containing lumbar regions L4-L6 was collected in an ice-cold cutting solution. Following sectioning, the slices were allowed to recover in warm artificial CSF (aCSF) containing (in mM): 25 D-glucose, 125 NaCl, 25 NaHCO_3_, 2.5 KCl, 1.25 NaH_2_PO_4_, 1 MgCl_2_, and 2 CaCl_2_, maintained at 34 °C for 30 min. All the solutions were continuously bubbled with 95% O_2_/5% CO_2_. After the initial recovery, the slices were left in aCSF for at least 60 min at room temperature (RT, ∼24 °) before transferring to the recording chamber for electrophysiological recordings. All recordings were carried out at RT.

### 2.3 Electrophysiology

For all the recordings, borosilicate glass pipettes (BF150-86-10, Sutter Instrument) with resistance 3-5 MΩ were pulled (P-1000, Sutter Instruments). The SDH neurons from laminae I-II were visualized using infrared differential interference contrast (IR-DIC) microscopy. tdTomato^+^ neurons of the SDH were identified using an LED illuminator (coolLed pE excitation system) excited at 565 nm with an appropriate filter (FITC) to visualize the tdTomato fluorescence. The neurons were patched to reach a seal resistance of >1.5 GΩ, and the seal was ruptured to attain whole-cell mode. We waited at least 5 min for proper diffusion of the internal solution throughout the cell before recording. The voltage-clamp recordings were acquired in gap-free mode, sampled at 20 kHz, and filtered at 2 kHz. The current-clamp recordings were acquired in episodic stimulation mode, sampled at a rate of 50 kHz, and filtered at 10 kHz.

All recordings were performed using a Multiclamp 700B amplifier (Molecular Devices) and were digitized by Digidata 1440 (Molecular Devices). Membrane potentials were not corrected for liquid junction potential. Series resistance was monitored continuously during all the recordings, and neurons with series resistance >15 MΩ and/or unstable baseline were excluded from the analysis.

#### 2.3.1 Current-clamp recordings

Current-clamp recordings were performed to validate the firing properties of tdTomato^+^ and tdTomato^−^ interneurons at the SDH of adult mice. To assess the firing properties of each neuron, steady hyperpolarizing and depolarizing current steps from −100 pA to 200 pA with increments of 50 pA were injected into the soma for a period of 500 ms or 1 s. All the recordings were carried out in neurons held at −60 mV by injecting the corresponding current. Potassium gluconate-based intracellular solution containing (in mM): 135 C_6_H_11_KO_7_, 6 NaCl, 2 MgCl_2_, 10 HEPES, 0.2 EGTA, 2 MgATP, and 0.3 NaGTP, pH adjusted to 7.25 with KOH and 280 mOsm, was used for all current-clamp recordings.

#### 2.3.2 Voltage-clamp Recordings

Voltage-clamp recordings were carried out to study the spontaneous inhibitory postsynaptic currents (sIPSCs) in both tdTomato^+^ and tdTomato^−^ interneurons of the SDH of adult and aged mice. All sIPSCs were recorded at −70 mV for 120 s using a Cs-based intracellular solution containing (in mM): 140 CsCl, 10 EGTA, 10 HEPES, 2 MgCl_2_, 2 MgATP, 4 Na_2_ATP, 0.4 Na_2_GTP, and 5 QX-314, pH adjusted to 7.3 with CsOH and 292 mOsm. This intracellular solution yielded a chloride reversal potential of ∼2 mV. The aCSF included 2,3-Dihydroxy-6-nitro-7-sulfamoyl-benzo(f)quinoxaline (NBQX, Alomone Labs) 20 μM and 2-amino-5-phosphonopentanoate (D-AP5, Abcam) 50 μM to block glutamatergic AMPA and NMDA currents, respectively.

### 2.4 Data analysis

The firing patterns of tdTomato^+^ and tdTomato^−^ neurons were analyzed and divided into tonic, delayed, initial burst, and single spike firing based on previous studies [1,2]. All sIPSC events were detected offline using the template search feature for event detection in ClampFit software (pClamp10, Molecular Devices). All sIPSC events <6 pA in amplitude and their corresponding inter-event intervals were discarded from the analysis. All the data were analyzed using GraphPad Prism 6th edition software (GraphPad Software, La Jolla, CA). Shapiro-Wilk test was used to determine the normality of the data. The significance in the differences of cumulative distribution curves of sIPSC amplitudes and inter-event intervals were analyzed using Kolmogorov-Smirnov (KS) non-parametric test. The comparison of the average amplitude and frequency of the sIPSCs was carried out using Mann-Whitney non-parametric test. For all tests, * *p* < 0.05 (two-sided) was considered significant. The individual values, obtained from a single cell, are represented as scattered dot plots with bar graphs of mean ± standard error of the mean (SEM). The data from the individual neurons are presented in the figures for all experiments. The description of the number of recorded cells (n), the number of slices per animal, the number of animals (N), and the specific statistical test used are detailed in the figure legends and in the main text.

### 2.5 Data availability

All datasets generated and/or analyzed during the current study are available in the main text or upon request.

Further information and requests for resources and reagents should be directed to and will be fulfilled by the corresponding contact, Alexander Binshtok.

## 3. Results and Discussion

Until recently, the differentiation of SDH inhibitory and excitatory interneurons was based only on their firing pattern, morphology, and neurochemical properties [1]. Recently, transgenic mice with fluorescently labeled inhibitory (*Vgat*) or excitatory (*Vglut*) spinal cord interneurons were developed [10]. Here we used *Vgat;tdTomato* transgenic line expressing tdTomato, specifically in the inhibitory neurons of the spinal cord (Figure 1A). It was previously shown that in *Vgat;tdTomato* mice, >90% of the immunolabeled tdTomato-expressing neurons are inhibitory, while the unlabeled cells are excitatory [10]. To confirm these results, we characterized the firing pattern of tdTomato^+^ and tdTomato^−^ neurons using spinal cord slices from adult *Vgat;tdTomato* mice. Earlier electrophysiological studies in the SDH showed that the majority of inhibitory neurons are tonically firing, i.e., generating multiple action potentials (APs) following near-threshold depolarizations [10,11]. The excitatory neurons are mostly delayed-firing, i.e., the first AP occurs with a prominent delay, with some exhibiting initial bursting and single AP firing patterns [11–13]. Consistently, we observed that 90% of the tdTomato^+^ interneurons exhibited a tonic firing pattern (9 of 10 neurons) while the tdTomato^−^ neurons showed non-tonic firing patterns consisting of delayed firing (9 out of 12 neurons), initial bursting (1 out of 12), and single spiking (2 out of 12 neurons, Figure 1A). These results further validated that the expression of tdTomato is limited to the inhibitory interneurons in our transgenic line.

**Figure 1.**
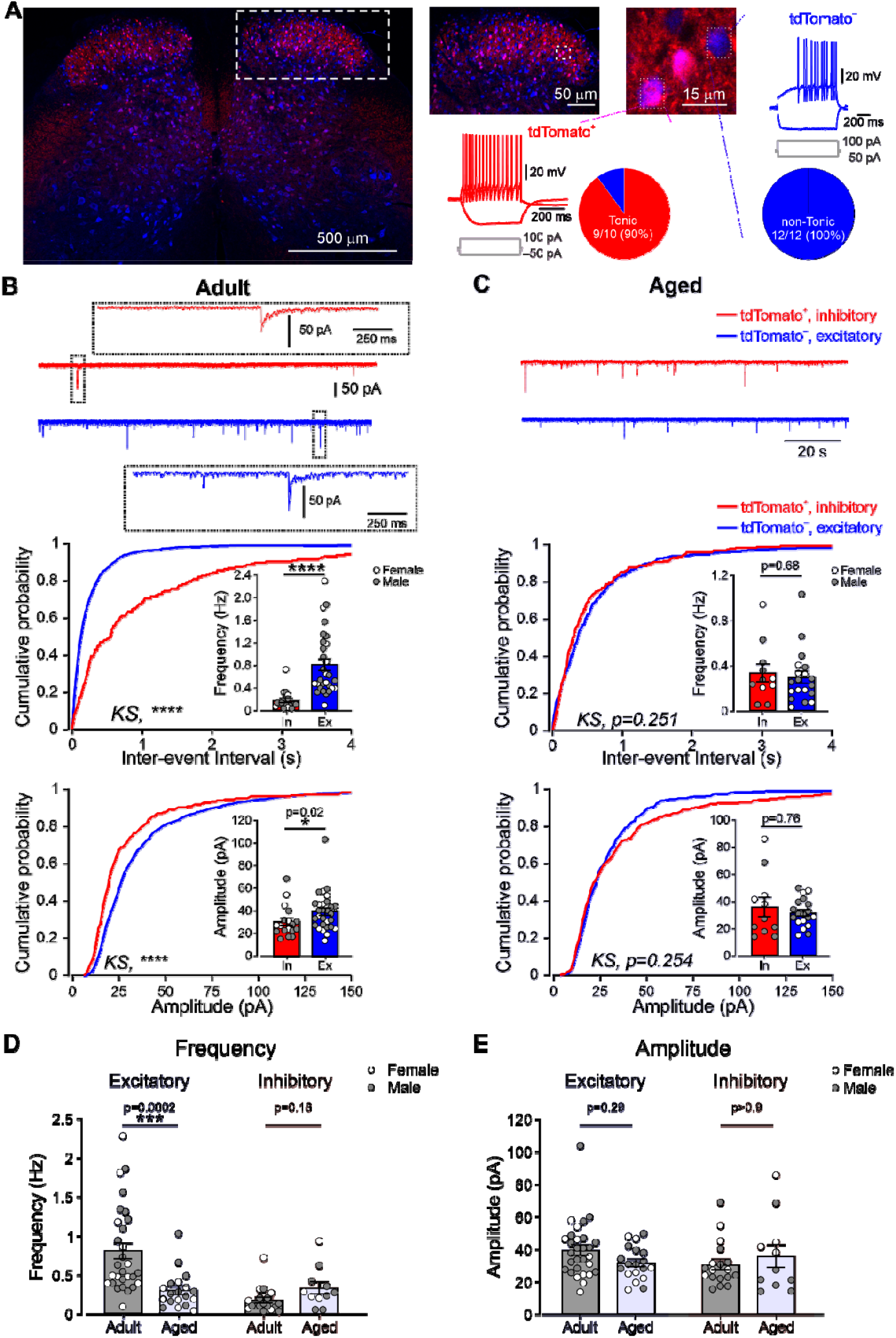
**A**. Representative image of a 30 μm-thick lumbar spinal cord (L4) section showing the distribution of tdTomato^+^ inhibitory (*purple*) and tdTomato^−^ excitatory interneurons (*blue*) in a spinal cord of *Vgat;tdTomato* transgenic mouse. Expanded images show regions of interest from the SDH with inhibitory interneurons and adjacent excitatory neurons. Representative of 6 mice. The color-coded insets show representative examples of the characteristic firing patterns of tdTomato^+^ and tdTomato^−^ interneurons from the SDH. For tdTomato^+^ neurons, representative of 9 out of 10 neurons. For tdTomato^−^ neurons, representative of 9 out of 12 neurons. The pie chart depicts the proportion of tonically firing neurons among all recorded tdTomato^+^ and tdTomato^−^ neurons. 9 of 10 neurons showed a tonic firing pattern and 1 delayed firing pattern. None of the 12 tdTomato^−^ neurons exhibited tonic firing patterns; 9 of 12 showed delayed firing, 2 of 12 fired a single spike, and 1 out of 12 showed initial bursting firing patterns. N = 3 mice for tdTomato^+^ and N = 6 mice for tdTomato^−^ neurons, 1-3 slices per mouse. **B**. *Upper*, Representative voltage-clamp traces of sIPSCs recorded from SDH inhibitory (*red*) and excitatory (*blue*) interneurons from adult mice. All recordings were carried out by holding the cell at −70 mV in the presence of 20 μM NBQX and 50 μM D-AP5 in the bath solution. Representative of 18 inhibitory and 29 excitatory neurons from 12 and 15 mice, respectively, 1-3 slices per animal. *Middle and Lower*, the cumulative distribution of inter-event intervals (*middle panel*) and amplitudes (*lower panel*) of sIPSCs in inhibitory (*red*) and excitatory (*blue*) neurons. Insets: Bar graphs and individual values (*white dots* for females and *grey dots* for males) comparing frequencies (*middle panel*) and amplitudes (*lower panel*) of sIPSCs in inhibitory (*red bar*) and excitatory (*blue bar*) neurons. Note that the frequency and amplitude of sIPSCs recorded from excitatory neurons are significantly higher than in inhibitory neurons. The Kolmogorov-Smirnov test was used to compare cumulative distributions between excitatory and inhibitory neurons. The Mann-Whitney test was used to compare frequencies and amplitudes between excitatory and inhibitory neurons. Inhibitory neurons: n = 18 neurons, N = 12 mice, 1-3 slices per animal. Excitatory neurons: n = 29 neurons, N = 15 mice, 1-3 slices per animal. **C**. Same as B, but for sIPSCs assessed in aged mice. The traces are representative of 11 inhibitory and 19 excitatory interneurons from 7 and 8 mice, respectively, 1-3 slices per animal. Note, no difference in frequency and amplitude of sIPSCs between excitatory and inhibitory neurons. The Kolmogorov-Smirnov (KS) test was used to compare cumulative distributions between excitatory and inhibitory neurons. The Mann-Whitney test was used to compare frequencies and amplitudes between excitatory and inhibitory neurons. Inhibitory neurons: n = 11 neurons, N = 7 mice, 1-3 slices per animal. Excitatory neurons: n = 19 neurons, N = 8 mice, 1-3 slices per animal. **D**. Bar graph (mean ± SEM) and individual values (*white dots* for females and *grey dots* for males) comparing sIPSC frequencies between excitatory and inhibitory neurons of adult and aged mice. Adult_Excitatory_: n = 29 neurons, N = 15 mice, 1-3 slices per animal; Aged_Excitatory_: n = 19 neurons, N = 8 mice, 1-3 slices per animal; Adult_Inhibitory_: n = 18 neurons, N = 12 mice, 1-3 slices per animal; Aged_Inhibitory_: n = 11 neurons, N = 7 mice, 1-3 slices per animal Kruskal-Wallis test with posthoc Dunn’s test. **E**. Same as D but comparing sIPSC amplitudes between excitatory and inhibitory neurons of adult and aged mice. Adult_Excitatory_: n = 29 neurons, N = 15 mice, 1-3 slices per animal; Aged_Excitatory_: n = 19 neurons, N = 8 mice, 1-3 slices per animal; Adult_Inhibitory_: n = 18 neurons, N = 12 mice, 1-3 slices per animal; Aged_Inhibitory_: n = 11 neurons, N = 7 mice, 1-3 slices per animal. Kruskal-Wallis test with posthoc Dunn’s test.

We next studied whether the inhibitory drive to the excitatory and inhibitory SDH interneurons alters with age by performing whole-cell recordings in voltage-clamp configuration from identified tdTomato^+^ inhibitory and tdTomato^−^ excitatory interneurons from the SDH of adult (3-5 m.o.) and aged (12-13 m.o.) *Vgat;tdTomato* mice. We show that in adult mice, the frequency of sIPSCs in inhibitory neurons was as low as 0.2 Hz (0.17±0.04 Hz, n = 18 neurons, N = 12 mice, Figure 1B). Notably, the sIPSC frequency in excitatory neurons was significantly higher (0.8±0.1 Hz, n = 29 neurons, N = 15 mice, p<0.0001, Mann-Whitney test, Figure 1B). At this age, the amplitude of sIPSCs in excitatory neurons was also significantly higher than in inhibitory SDH neurons (Figure 1B).

Importantly, in aged mice (12-13 m.o.), at the time point in which the increased responsiveness to noxious stimuli has been reported [14], the frequencies and amplitudes of the sIPSCs were indistinguishable between excitatory and inhibitory interneurons (Figure 1C).

These data suggest that in adult mice, excitatory neurons receive stronger inhibitory input than inhibitory interneurons. Considering that excitatory neurons comprise ∼70% of the neurons within SDH [1], the relative increase in the inhibitory synaptic transmission onto the excitatory neurons might help maintain the excitation-inhibition equilibrium of the SDH network. Notably, we show that this compensation disappears in aged animals.

Furthermore, we show that the frequency of sIPSCs on excitatory but not inhibitory neurons was significantly lower in aged mice than in adult mice (Figure 1D). We did not observe any differences in the amplitudes of sIPSCs (Figure 1E).

Altogether, our results demonstrate that (1) in the adult mice, the inhibitory transmission onto excitatory SDH neurons is higher than on the inhibitory SDH interneurons, and (2) the inhibitory transmission onto excitatory SDH neurons decreased with age. The net effect of the age-dependent decreased inhibition of excitatory neurons might lead to overall excitation of the SDH network, plausibly contributing to increased pain.

Our data showing a decrease in the frequency of sIPSCs on excitatory neurons in aged mice implies that the age-related abolition of the differences in the inhibitory drive between excitatory and inhibitory neurons (see Figure 1C) could result from a decrease in inhibitory drive on excitatory neurons.

Our study has limitations as it does not fully explore the mechanisms behind the age-mediated alterations in synaptic transmission. The changes in frequency but not in the amplitude of sIPSCs with age could point to a presynaptic mechanism. One option is an age-dependent decrease in the number of inhibitory neurons. If this is the case, we would expect a decrease in the synaptic inputs to both excitatory and inhibitory neurons. However, we did not observe any changes in the inhibitory drive to inhibitory neurons (Figure 1D and E). Alternatively, the decrease in the inhibitory drive on the excitatory neurons could also result from decreased firing of presynaptic inhibitory neurons. This is also less likely because a recent study demonstrated no change in the firing frequency of inhibitory SDH neurons in aged mice [15]. Future experiments assessing changes in a spontaneous release (by measuring miniature IPSCs), examining the changes in the number of inhibitory terminals on excitatory neurons and the release probability from these synapses could provide further insight into the mechanism of age-dependent disinhibition on excitatory SDH neurons.

The age-mediated changes in synaptic connectivity in the SDH were previously examined [15]. However, unlike our results, the authors did not demonstrate any alterations in the frequencies of the sIPSCs. This discrepancy could be explained by the fact that Mayhew et al. did not distinguish between sIPSCs recorded from inhibitory and excitatory interneurons and pooled together data from all recorded SDH neurons. It is, therefore, reasonable that the averaging between excitatory and inhibitory neurons could have masked the differences in sIPSCs and their changes with age, as we observed in this study.

Taken together, the reduction in the inhibitory synaptic drive onto the excitatory interneurons in the SDH of aged mice might lead to overall disinhibition and, thus, hyperexcitability of the SDH network. In addition to the immediate impact on pain sensation, these age-mediated hyperexcitability changes might induce sensitization of the SDH network, increasing the likelihood of the transition from normal to pathological chronic pain in the elderly.

## Declarations

### Ethics approval

All experiments were conducted in accordance with the guidelines and approval of the animal ethics committee of the Hebrew University of Jerusalem (MD18-15604).

## Authors’ contributions

*Conceptualization*, P.R.R. and A.M.B.; *Methodology*, P.R.R, S.L. and A.M.B.; *Investigation*, P.R.R. and A.M.B.; Writing - Original Draft, P.R.R, S.L. and A.M.B.; Funding Acquisition, A.M.B.; Supervision, A.M.B.

## List of abbreviations

aCSF: artificial cerebrospinal fluid
D-AP5: 2-amino-5-phosphonopentanoate
GABA: γ-aminobutyric acid
IR-DIC: infrared differential interference contrast
KS: Kolmogorov-Smirnov
NBQX: 2,3-Dihydroxy-6-nitro-7-sulfamoyl-benzo(f)quinoxaline
RT: room temperature
SDH: superficial spinal dorsal horn
sEPSC: spontaneous excitatory postsynaptic currents
sIPSC: spontaneous inhibitory postsynaptic currents
Vgat: vesicular GABA transporter

